# Measuring the quality of scientific references in Wikipedia: an analysis of more than 115M citations to over 800,000 scientific articles

**DOI:** 10.1101/2020.04.08.031765

**Authors:** Joshua M. Nicholson, Ashish Uppala, Matthias Sieber, Peter Grabitz, Milo Mordaunt, Sean Rife

**Author notes:** Address correspondence to: Joshua M. Nicholson, PhD, scite Inc., 2 N 6th St, #31A, Brooklyn, NY 11249, USA.

## Abstract

Wikipedia is a widely used online reference work which cites hundreds of thousands of scientific articles across its entries. The quality of these citations has not been previously measured, and such measurements have a bearing on the reliability and quality of the scientific portions of this reference work. Using a novel technique, a massive database of qualitatively described citations, and machine learning algorithms, we analyzed 1,923,575 Wikipedia articles which cited a total of 824,298 scientific articles, and found that most scientific articles (57%) are uncited or untested by subsequent studies, while the remainder show a wide variability in contradicting or supporting evidence (2-41%). Additionally, we analyzed 51,804,643 scientific articles from journals indexed in the Web of Science and found that most (85%) were uncited or untested by subsequent studies, while the remainder show a wide variability in contradicting or supporting evidence (1-14%).

---

Wikipedia, the free online encyclopedia, is an integral part of the web and society. With over 18 billion visits per month, currently ranking it as the 10th most visited website in the world, it has become the go-to source of information for nearly all aspects of life. It is comprised of over 6M articles and 49M pages, which have received 934M edits from 38M users. Because Wikipedia is so important for maintaining a well-informed society, we sought to determine how primary research articles informing Wikipedia articles have been cited within the scientific community.

As of 2018, there were 824,298 scientific articles in Wikipedia referenced across 1,923,575 Wikipedia articles, meaning 32% (1,923,575/6,006,758) of all Wikipedia articles reference a scientific article. The accuracy of these articles is paramount, especially considering that Wikipedia is often the first and only source of information for some readers. The task is delegated to its large community of volunteer editors and users; claims are heavily debated and calls for primary sources of evidence are flagged with the now popular phrase: “Citation Needed.” But just how reliable are these sources?

To answer this question, we performed a citation analysis of scientific articles referenced in Wikipedia using “Smart Citation” data from scite. Smart citations provide the context for each citation and a classification describing whether it provides supporting or contradicting evidence for the cited claim. Classifications are performed by a deep learning model that has been trained on 43,665 expert-labeled citation statements with precision scores of 0.800, 0.8519, and 0.9615 for supporting, contradicting, and mentioning classifications, respectively (internal scite benchmarking data). To date, scite has analyzed over 15M full-text scientific articles, extracting over 500M citation statements that cite over 34M articles. These scientific articles were obtained through a variety of means, including retrieval of open access papers, preprints, PubMed Central, and through partnerships with various publishers.

Using this information, we analyzed the 824,298 scientific articles referenced in Wikipedia to see how they had been cited in the scientific literature. These articles have received 115,046,571 total Smart Citations according to scite. Of those Smart Citations, 3,435,635 (2.99%) indicate that they provide supporting evidence, 401,472 (.35%) indicate that they provide contradicting evidence, and 111,209,464 (96.7%) mention the citing study without indicating that they provide supporting or contradicting evidence. Wikipedia articles referencing scientific articles cited 2.44 (SD = 24.09) scientific articles on average. This figure differs slightly from a recent estimate likely due to variations in data collection (Arroyo-Machado et al. utilized Altmetric data in their analyses, while we used data retrieved directly from Wikipedia) [1]. Among scientific articles referenced by Wikipedia articles, the average number of citations was 130.52 (SD=476.93), the mean number of supporting citations was 3.96 (SD=9.98), the mean number of contradicting citations was. 45 (SD=1.34), and the mean number of mentioning citations was 126.10 (SD=470.90) (Table 1). The most cited scientific article referenced in Wikipedia describes Laemmli buffer, which is widely used in protein analysis and has over 66k citation statements [2]. Most articles (324,247/824,298, 39.34%) remain untested by other subsequent citing articles (mentioning cites with no supporting or contradicting cites), 148,243 (17.98%) have no citations at all, 235,036 (28.51%) have been supported with no contradicting evidence, 102,719 (12.46%) have been disputed with both supporting and contradicting evidence, and 14,052 (1.70%) have been contradicted with no supporting cites (Figure 1). 297 scientific papers referenced by Wikipedia articles have been retracted; however, the most of these references are recognized as retracted in the text of the Wikipedia article itself (for example, the Wakefield et al. [3] paper presenting evidence of a causal link between vaccines and autism is frequently cited as part of a discredited body of research).

**Table 1.** Citation Breakdown of Scientific Articles Referenced in Wikipedia

| **Classification** | **Wikipedia**<br>**Mean (SD)** | **Web of Science**<br>**Mean (SD)** |
| --- | --- | --- |
| Total citations | 130.52 (476.93) | 16.91 (58.20) |
| Mentioning citations | 126.10 (470.90) | 16.06 (56.72) |
| Supporting citations | 3.96 (9.98) | 0.75 (2.22) |
| Contradicting citations | 0.45 (1.34) | 0.11 (0.46) |

**Figure 1.**
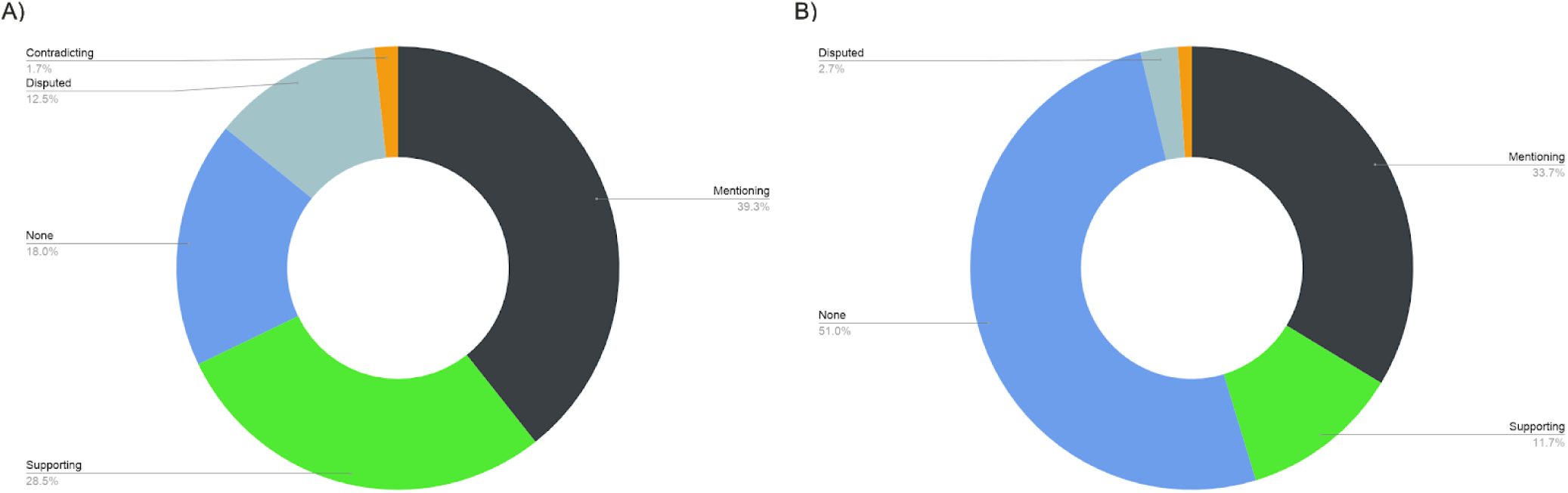
Overview of how scientific articles referenced in Wikipedia and articles from journals indexed in the Web of Science have been cited within the scientific literature. A) Most scientific articles referenced in Wikipedia (324,247/824,298, 39.34%) remain untested by other subsequent citing articles (mentioning citations with no supporting or contradicting citations), 148,243 (17.98%) have no citations at all, 235,036 (28.51%) have been supported with no contradicting evidence, 102,719 (12.46%) have been disputed with both supporting and contradicting evidence, and 14,052 (1.70%) have been contradicted with no supporting cites. B) Most scientific articles in Web of Science 17,441,574 (33.67%) have not been tested by other subsequent citing articles (has mentioning cites only) 26,396,010 (50.96%) has no citations at all, 6,038,194 (11.66%) have been supported with no contradictions, 1,407,829 (2.72%) have been disputed with both supporting and contradicting evidence, and 521,024 (1.01%) have been contradicted with no supporting cites.

To see how scientific articles referenced in Wikipedia compare to the scientific literature as a whole, we looked at the citation breakdown of 51,804,643 articles (429,780,086 total Smart Citations) from journals indexed in the Web of Science. Of these citations, 18,940,149 (4.41%) indicate that they provide supporting evidence, 2,710,605 (.63%) indicate that they provide contradicting evidence, and 408,129,332 (94.96%) mention the citing study without indicating that they provide supporting or contradicting evidence. Similar to Wikipedia, most articles 17,441,574 (33.67%) have not been tested by other subsequent citing articles (has mentioning cites only), 26,396,010 (50.96%) has no citations at all, 6,038,194 (11.66%) have been supported with no contradictions, 1,407,829 (2.72%) have been disputed with both supporting and contradicting evidence, and 521,024 (1.01%) have been contradicted with no supporting cites. The average number of citations articles from these journals received was 16.91 (SD=58.20), the mean number of supporting citations was. 75 (SD=2.22), the mean number of contradicting citations was. 11 (SD=.46), and the mean number of mentioning citations was 16.06 (SD=56.71) (Table 1).

Our results should be considered with caution given the limitations of the model precision, the current limited coverage of articles analyzed by scite, and that articles without DOIs or identifiable DOIs in the data set were excluded. Beyond technical limitations, it is also important to consider what the citation classifications mean. For example, a contradicting citation statement does not necessarily mean the cited paper is wrong because: 1) scite classifies citation statements at the level of the claim, not the full paper, and 2) the citing article making the contradicting claim itself could be without merit. Nonetheless, these numbers are a good approximation of how the scientific foundations of Wikipedia have been tested in the scientific literature and represent the first time an analysis of the quality of citations, not just the quantity, has been done at this scale. Previous citation analyses at the individual article level have shown that reporting the citation context can be informative for readers [4][5] with one citation analysis [5] causing the publisher to add the following warning to the original report [6], “*Editor’s Note (added May 31, 2017): For reasons of public health, readers should be aware that this letter has been “heavily and uncritically cited” as evidence that addiction is rare with opioid therapy.”*

To look at how citation context could impact Wikipedia users if it were linked next to scientific references, we examined a handful of articles directly. The Wikipedia article on “Amygdala” states, “*In 2006, researchers observed hyperactivity in the amygdala when patients were shown threatening faces or confronted with frightening situations. Patients with severe social phobia showed a correlation with increased response in the amygdala”* citing Phan et al. [7] as evidence for this statement. According to scite [8], this reference has received 259 mentioning citation statements, 23 supporting citation statements, and 3 contradicting citation statements (Figure 2). Thus, while some have provided supporting evidence, two studies have called this into question, with one report stating [9], “*These findings do not replicate previous studies...”* The citation context offers a more complete picture, potentially affecting decisions by everyday readers and choices of editors. Consider the Wikipedia article “Suicide and Internet” which features the following statement, “*A survey has found that suicide-risk individuals who went online for suicide-related purposes, compared with online users who did not, reported greater suicide-risk symptoms, were less likely to seek help and perceived less social support,”* highlighting a report by Harris, McLean, and Sheffield [10]. As identified by scite [11], this report was later contradicted by a subsequent study finding that suicide-related Internet use individuals were more likely to seek help [12] (Figure 3).

**Figure 2.**
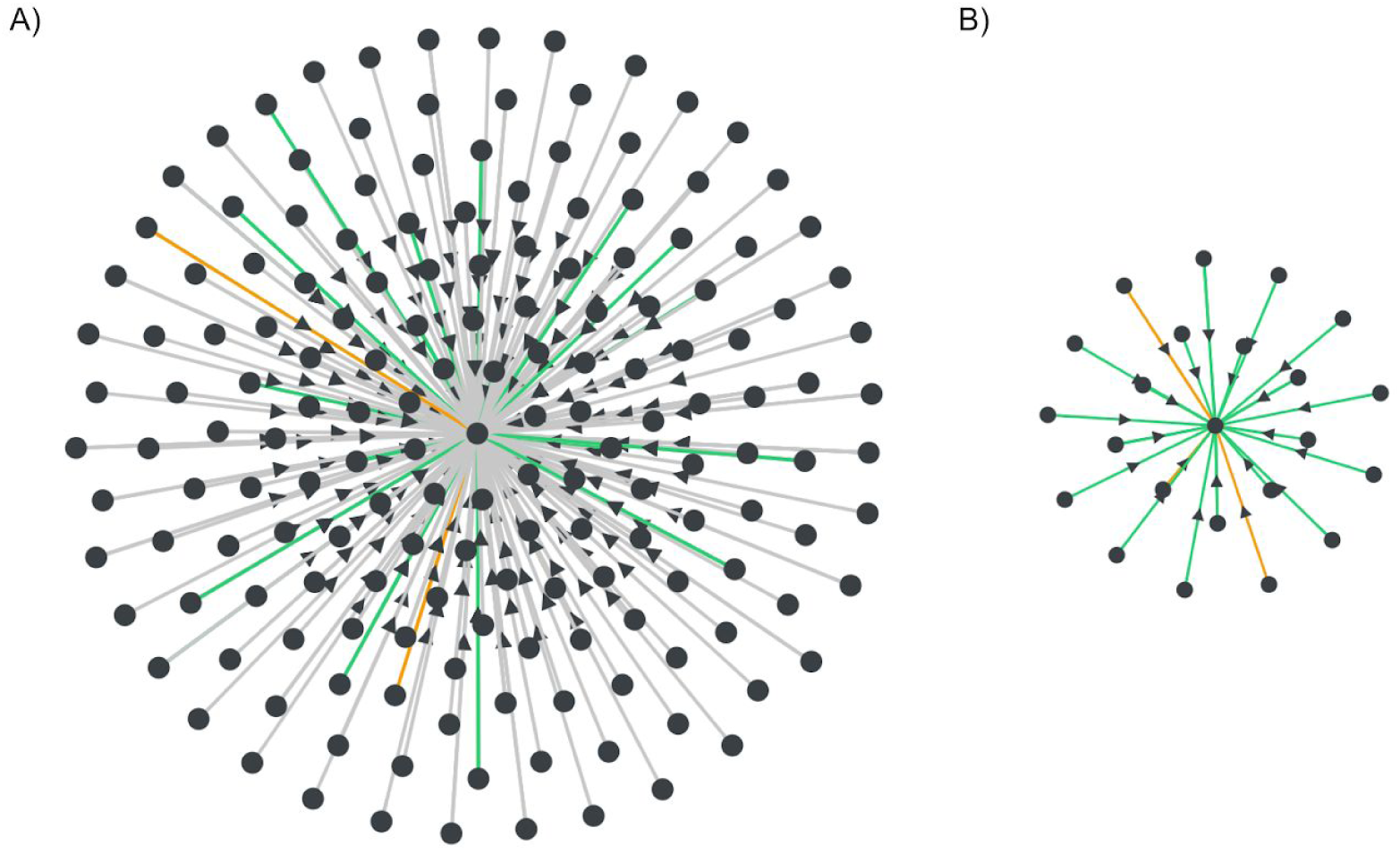
Citation visualization of Phan et. al [7]. A) Citation network showing supporting (green), contradicting (orange), and mentioning (grey) citations. B) Citation network showing only supporting and contradicting cites. Interactive figure can be accessed here: https://scite.ai/visualizations/association-between-amygdala-hyperactivity-to-gVamGz

**Figure 3.**
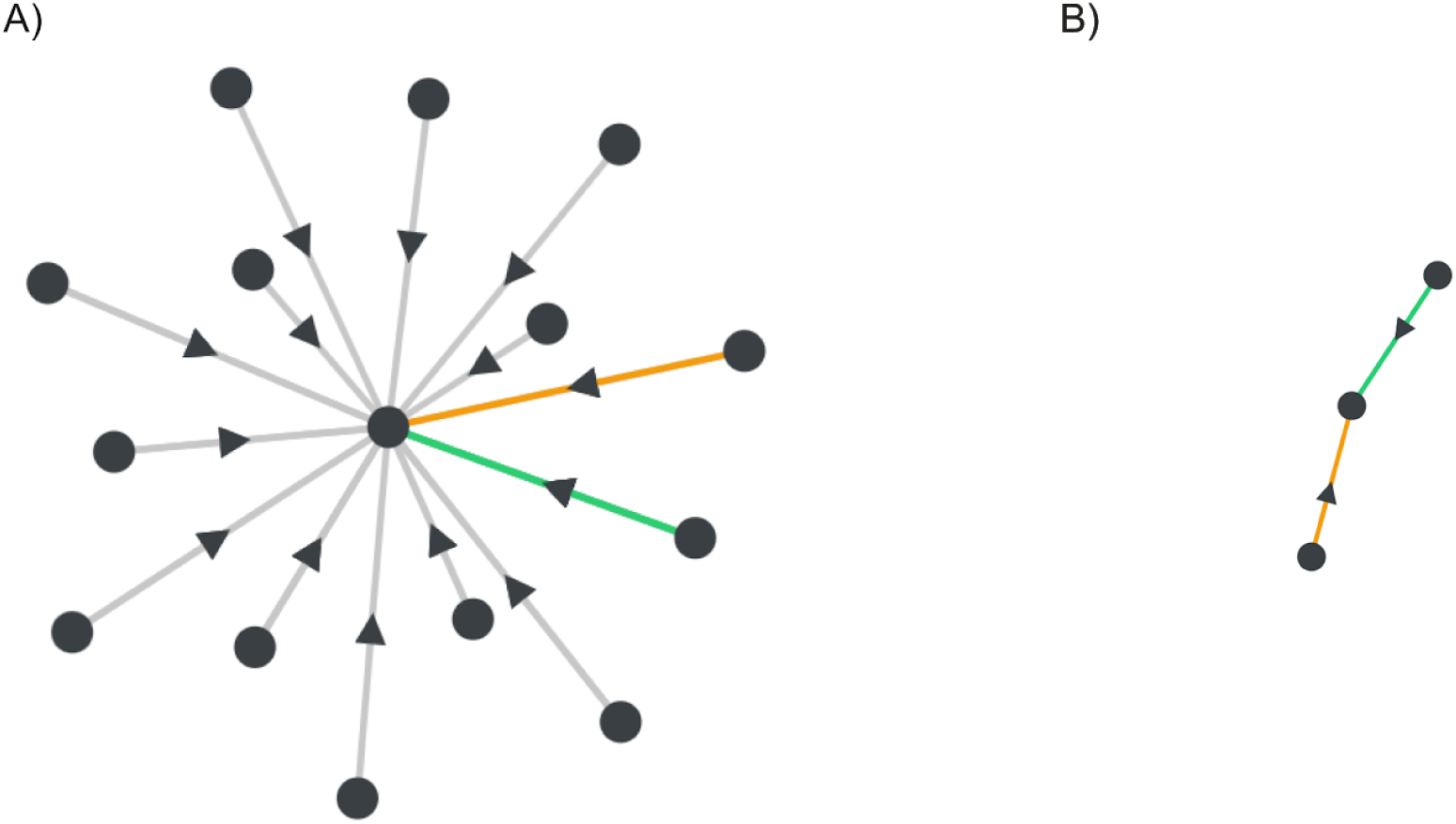
Citation visualization of Harris, McLean, and Sheffield [10]. A) Citation network showing supporting (green), contradicting (orange), and mentioning (grey) citations. B) Citation network showing only supporting and contradicting cites. Interactive figure can be accessed here: https://scite.ai/visualizations/examining-suicide-risk-individuals-who-go-RyyzV5

Another study [13] does provide supporting evidence noting, “*Our findings supported previous research showing suicide risk was related to greater likelihood of online interpersonal communications,”* however the supporting paper is from the same group as the original paper while the contradicting citation comes from an independent group while. Thus, providing contextual citation information for this Wikipedia claim could influence behavioral choices that have potentially life or death consequences for a large population of people.

In conclusion, Wikipedia largely references scientific articles that are supported or untested with a minority being contradicted, similar to what has been seen in other citation network analyses and similar to the scientific literature as a whole [14]. When Wikipedia articles cite scientific papers that have been subsequently retracted, this is often explicitly stated, and often in service of a larger conversation about the article itself. However, citation numbers alone fail to capture the tenuous nature of scientific claims. Making the citation context available to moderators and readers is critical to reliably evaluating scientific claims and we suggest the adage “Citation Needed” is not enough. References in Wikipedia as well as scientific articles themselves should display citation contexts. Platforms and publishers like <u>**Europe PMC**</u> and <u>**Wiley**</u> are starting to adopt this approach and technology and we think this could be helpful for Wikipedia as well.

## Methods Supplement

### Identification research articles in Wikipedia

We used data previously scraped from Wikipedia [15] containing a list of citations with their identifiers from Wikipedia content dumps published on March 1, 2018. The data fields included the *id* and *type*, such as “pmid” for PubMed ID, “pmcid” for PubMed Central ID, and “doi” for Digital Object Identifier. First, we mapped all identifiers to DOIs using mapping data from the PMC metadata database (https://www.ncbi.nlm.nih.gov/pmc/pmctopmid/), which provides links between PMIDs, PMCIDs, and DOIs. Mapped DOIs were combined with DOIs where the identifier type was designated “doi” and were considered valid if the DOI existed in a dataset of all known DOIs provided by CrossRef. Within the scraped data, 96% of entries were successfully linked to a DOI, and among those DOIs, 98% were valid. Given a valid DOI, it was possible to query against our internal citation data to determine how frequently it was cited, supported, mentioned, or contradicted.

### Citation analysis

Citation analyses were performed by querying internal scite citation data. Descriptive analyses and graph generation were performed in R. All queries and code can be found at https://github.com/scitedotai/research-wikipedia.

## Notes

**Conflicts of Interest** The authors are shareholders and/or consultants or employees of Scite Inc.

https://github.com/scitedotai/research-wikipedia

### Competing Interest Statement

The authors are shareholders and/or consultants or employees of Scite Inc.

